# GUN1-independent retrograde signaling targets the ethylene pathway to repress photomorphogenesis

**DOI:** 10.1101/2020.09.02.280206

**Authors:** Charlotte M. M. Gommers, María Águila Ruiz-Sola, Alba Ayats, Lara Pereira, Marta Pujol, Elena Monte

**Affiliations:** Plant Development and Signal Transduction Program, Center for Research in Agricultural Genomics (CSIC- IRTA-UAB-UB), Barcelona, Spain; Laboratory of Plant Physiology, Wageningen University & Research, Wageningen, The Netherlands; Plant and Animal Genomics Program, Center for Research in Agricultural Genomics (CSIC- IRTA-UAB-UB), Barcelona, Spain; Institut de Recerca i Tecnologia Agroalimentàries (IRTA), Barcelona, Spain; Consejo Superior de Investigaciones Científicas (CSIC), Barcelona, Spain; Center for Applied Genetic Technologies, University of Georgia, Athens, United States

**Author notes:** Author for contact: Elena Monte. Author contributions: C.G. and E.M. conceived the project. C.G., M.R-S., A.A., L.P. and M.P. planned and performed the experiments. C.G., M.R-S. and A.A. analyzed the data. C.G. and E.M. wrote the manuscript.

## Abstract

When germinating in the light, *Arabidopsis* seedlings undergo photomorphogenic development, characterized by short hypocotyls, greening and expanded cotyledons. Stressed chloroplasts emit retrograde signals to the nucleus that induce developmental responses and repress photomorphogenesis. The nuclear targets of these retrograde signals are not yet fully known. Here, we show that lincomycin-treated seedlings (which lack developed chloroplasts) show strong phenotypic similarities to seedlings treated with ethylene (ET) precursor 1-aminocyclopropane-1-carboxylic acid (ACC), as both signals inhibit cotyledon separation in the light. We show that the lincomycin-induced phenotype partly requires a functioning ET signaling pathway, but could not detect increased ET emissions in response to lincomycin treatment. The two treatments show overlap in up-regulated gene transcripts, downstream of transcription factors ETHYLENE INSENSITIVE3 (EIN3) and EIN3-LIKE1 (EIL1). The induction of the ethylene signaling pathway is triggered by an unknown retrograde signal acting independently of GENOMES UNCOUPLED1 (GUN1). Our data show how two apparently different stress responses converge to optimize photomorphogenesis.

**One Sentence Summary:** Chloroplast retrograde signaling targets the ethylene-regulated gene network to repress photomorphogenesis in Arabidopsis

## Introduction

As photoautotrophs, plants depend on light for growth and survival. Chloroplasts absorb photons to fuel the photosynthesis reaction, which is the basis of all carbon sources on earth. Most chloroplast proteins are encoded in the nuclear genome, so tight communication between chloroplasts and nucleus is essential to build and maintain the photosynthesis apparatus and to accurately adjust to environmental stresses with potential damage to the photosystems. Retrograde signals from the plastid inform the nucleus about disruptions in its function or development (reviewed in (Chan et al., 2016; Crawford et al., 2018).

Quickly after germination, light triggers a developmental program which includes the opening and expansion of cotyledons, chloroplast maturation, reduced hypocotyl elongation and promoted root growth, called photomorphogenesis (Gommers and Monte, 2018). The initiation of photomorphogenesis depends on the activation of a set of photoreceptors, sensitive to red and far-red (phytochromes, phys), or blue light (cryptochromes, crys). These repress a set of transcription factors, including PHYTOCHROME INTERACTING FACTORS (PIFs) and ETHYLENE INSENSITIVE3 (EIN3), that are repressors of photomorphogenesis in darkness (Leivar et al., 2008; Shi et al., 2018).

When chloroplast development is interrupted, by chemicals such as lincomycin (inhibitor of chloroplast translation) or norflurazon (inhibitor of carotenoid synthesis), or by high light, this induces a retrograde signaling (RS) cascade and inhibits cotyledon separation, possibly to minimize the stress-exposed area and protect the apical meristem (Ruckle and Larkin, 2009; Martín et al., 2016). This RS pathway acts via the chloroplast localized, nuclear encoded, protein GENOMES UNCOUPLED1 (GUN1), a master integrator of RS. GUN1 is a plastid-localized member of the pentatricopeptide repeat family (PRR) involved in chloroplast synthesis as well as stress signaling, especially early during leaf development (Koussevitzky et al., 2007; Jia et al., 2019; Pesaresi and Kim, 2019). The lincomycin-induced, GUN1-mediated, RS pathway antagonizes phytochrome signaling and targets genes which encode proteins that promote photomorphogenesis and are repressed by PIFs in darkness. One of such genes encodes the transcription factor GOLDEN2 LIKE1 (GLK1). *GLK1* transcriptional repression by GUN1-mediated RS contributes to the closed cotyledon-phenotype in lincomycin-treated seedlings (Martín et al., 2016). An additional RS pathway with direct effect on light signaling is mediated by the plastid stress molecule methylerythritol cyclodiphosphate (MEcPP), which promotes phyB accumulation and suppresses auxin and ethylene-mediated hypocotyl elongation in red light (Jiang et al., 2020).

Another stress signal that affects photomorphogenesis is accumulation of the gaseous phytohormone ethylene (ET). ET biosynthesis is induced during environmental stresses such as pathogen attack or vegetation shade, it accumulates in the plant in situations where the gas-flow is limited (flooding, under a pressing soil layer, or in close canopies), and acts as a neighbor detection molecule (Dubois et al., 2018). Depending on the concentration, light availability and species, ET can either suppress or induce plant growth (Pierik et al., 2006; Dubois et al., 2018). In young *Arabidopsis* seedlings, it represses several aspects of photomorphogenesis, such as hypocotyl growth arrest and cotyledon separation and expansion (Das et al., 2016; Shi et al., 2016; Shi et al., 2018).

Ethylene is perceived by multiple ethylene receptors located at the golgi and ER membranes: ETHYLENE RESPONSE SENSOR 1 (ERS1), ERS2, ETHYLENE RESISTANCE 1 (ETR1), ETR2 and ETHYLENE INSENSITIVE 4 (EIN4) (Lacey and Binder, 2014). In the absence of ET, these receptors are active and activate the kinase CONSTITUTIVE TRIPLE RESPONSE1 (CTR1), an inhibitor of the ER-localized, NRAMP-like protein ETHYLENE INSENSITIVE2 (EIN2). In the nucleus, EIN3-BINDING F-BOX1 (EBF1) and EBF2 bind and degrade the transcription factors EIN3 and EIN3-LIKE1 (EIL1). In the presence of ET, the receptors and therefore CTR1, are inactive. This renders active EIN2, that inhibits the translation of EBF1 and EBF2 mRNA. As a consequence, EIN3 and EIL1 accumulate and target ET-regulated genes to optimize development to the stress situation that caused ET to accumulate (Binder, 2020).

Even though lincomycin-induced RS and ET cause similar phenotypes in light-grown seedlings, with MEcPP affecting ET-mediated hypocotyl elongation (Vogel et al., 2014; Jiang et al., 2020), and high light has been shown to induce a stress-pathway via ETHYLENE RESPONSE FACTORS (ERFs) (Vogel et al., 2014), very little is known about a possible overlap between these stress signaling pathways. Here, we show that chloroplast RS promotes the expression of ET-responsive genes, which causes the delayed photomorphogenesis in light-grown plants. We present evidence that this RS-pathway acts independent of the well-studied GUN1-mediated signaling cascade. Our findings additionally suggest that various environmental stresses converge at a similar core-set of nuclear genes to change plant development and prevent further damage.

## Results

### Ethylene affects photomorphogenesis in continuous light

Previous work has shown that ethylene can affect seedling photomorphogenesis. Dark-grown *EIN3/pifqein3eil1* seedlings treated with ET precursor 1-aminocyclopropane-1-carboxylic acid (ACC) had small, partly unseparated cotyledons but short hypocotyls compared to seedlings grown on regular MS medium (Shi et al., 2018). Dark-grown *ein3eil1* seedlings transferred to light had larger and more separated cotyledons, while *EIN3-ox* displayed unopened and smaller cotyledons. In continuous red light, ACC triggered hypocotyl elongation, whilst inhibiting cotyledon expansion in wild type and *EIN3-ox*, but not in *ein3eil1* (Shi et al., 2016). Similar phenotypes were found when short day-grown wild-type seedlings were treated with ethylene gas (Das et al., 2016). To confirm the effect of ET on several aspects of photomorphogenic growth in continuous white light-grown *Arabidopsis thaliana* seedlings, we germinated seeds on media with different concentrations of ACC and analyzed hypocotyl length, cotyledon unfolding and apical hook opening. Compared to the control, low levels of ACC caused a slight increase of hypocotyl length in wild type (WT) seedlings. This effect was reversed by higher concentrations (> 5 µM), as visualized by the heatmap in figure 1A, which represents relative values compared to WT without ACC, and Fig. S1. At these higher concentrations, other aspects of photomorphogenesis were repressed, resulting in partly unseparated cotyledons and apical hook formation (Fig. 1A-C, S1). ACC additionally strongly inhibited primary root growth (Fig. 1B). All these phenotypes depended on the activation of the ethylene pathway via EIN2 and EIN3/EIL1, shown by the ACC-insensitivity of *ein2-5* and *ein3eil1* mutants. For further experiments we focused on the cotyledon phenotype using 10 µM ACC, a concentration that strongly prevented cotyledon separation in continuous light without compromising overall growth.

**Figure 1.**
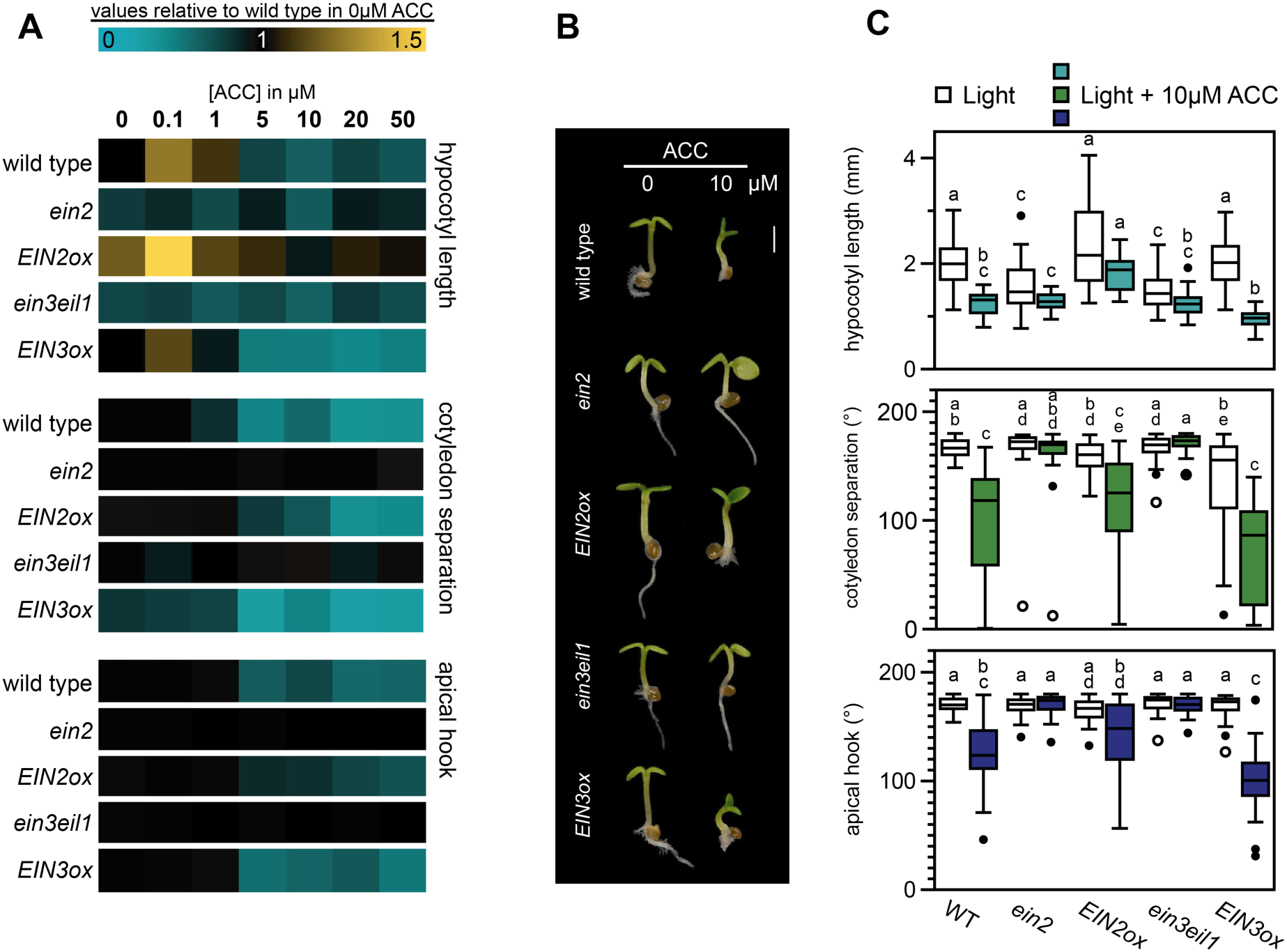
ACC inhibits photomorphogenesis in the light via EIN2 and EIN3. **A**. Relative hypocotyl length, cotyledon separation (presented as the angle between cotyledons) and apical hook angle of three-day old wild type Col-0 (WT), *ein2, 35S::EIN2-GFP/ein2* (*EIN2ox*), *ein3eil1* and *35S::EIN3-GFP/ein3eil1* (*EIN3ox*) seedlings grown in continuous low light (approx. 1.5 µmol m^-2^ s^-1^) on medium supplemented with different concentrations of ACC as compared to normal growth medium (complete dataset in Fig. S1). **B**. Representative seedlings grown in control medium or supplemented with 10 µM ACC. Scale bar = 1 mm. **C**. Boxplots representing the absolute data for control and 10 µM ACC-treated seedlings. Same data as in A. Different letters indicate significant differences, Kruskal-Wallis with post-hoc Dunn test. In A and C, biological replicates n = approx. 40 seedlings.

### Suppression of photomorphogenesis by retrograde signals requires the ethylene signaling pathway

The closed cotyledon phenotype caused by ACC resembles the phenotype seen in light-grown *Arabidopsis* seedlings treated with chloroplast inhibitor lincomycin (Fig. 2A) (Martín et al., 2016), suggesting that both might be regulated by a similar pathway. To test this possibility, we treated *ein2* and *ein3eil1* mutant seedlings with lincomycin. As shown in Fig. 2A, these ET mutants were partially insensitive to the lincomycin treatment and cotyledon separation was only partially inhibited compared to wild type. The WT-like phenotype of the single *ein3* and *eil1* mutants as compared to the significantly distinct cotyledon phenotype of *ein3eil1* in lincomycin, shows that EIN3 and EIL1 act redundant in the repression of cotyledon opening (Fig. S2A). Over-expression of EIN3 slightly, but significantly, enhanced the lincomycin-induced phenotype compared to the wild-type control. Contrastingly, the *ctr1* mutant, with a constitutive ET response, displayed constitutively closed cotyledons and was insensitive to lincomycin (Fig. 2A). Together, these results suggest that lincomycin-induced inhibition of cotyledon separation in light-grown seedlings requires ET signaling. These findings were further confirmed by chemical inhibition of ethylene perception by silver (AgNO_3_), which did not affect wild-type seedlings under normal light conditions but partially blocked the lincomycin-induced inhibition of cotyledon separation (Fig. 2B). The inhibition of cotyledon separation is less, but significantly, repressed by norflurazon-induced retrograde signaling, and could be compromised by inhibition of ET perception using AgNO_3_ as well (Fig. S2B).

**Figure 2.**
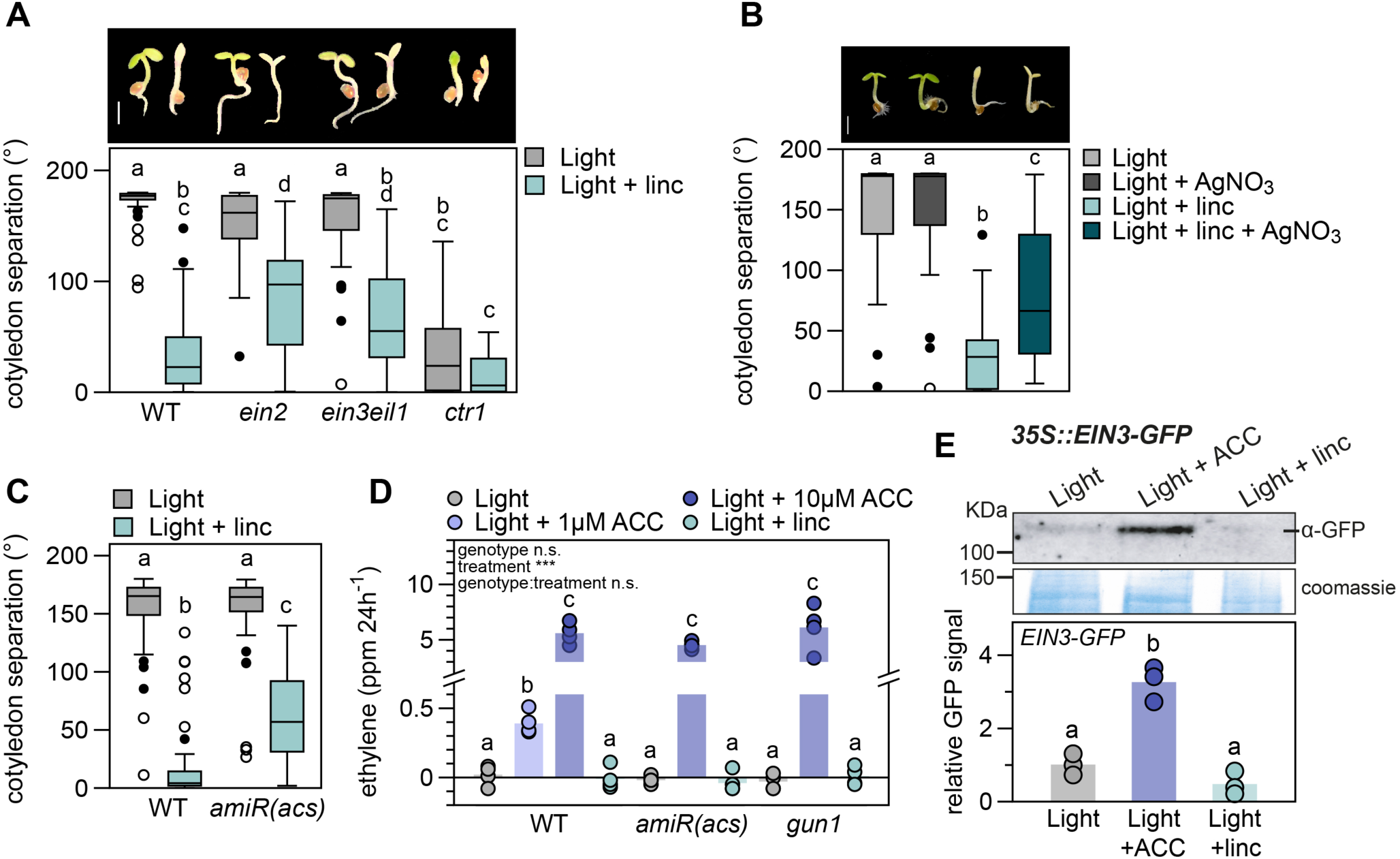
Lincomycin-mediated repression of cotyledon separation acts via the ET signaling pathway without altering ET emission and EIN3 levels. **A**. Cotyledon separation (degrees) of three-day-old, light-grown wild type (WT), *ein2, ein3eil1* and *ctr1* seedlings on regular growth medium or medium supplemented with 0.5 mM lincomycin (linc). Pictures are representative seedlings, scale bar = 1 mm. **B**. Cotyledon separation of three-day-old, light-grown wild type seedlings on medium supplemented with 5 µM AgNO_3_, 0.5 mM linc, or a combination of both. Pictures are representative seedlings, scale bar = 1 mm. **C**. Cotyledon separation of WT and ethylene deficient *amiR(acs)* mutants, grown as in A. For A – C, biological replicates n = approx. 40 seedlings. **D**. Ethylene emission (in ppm / 24 hours) of 120 WT, *amiR(acs)* and *gun1* seedlings grown in light, on regular growth medium, or supplemented with 1 µM (WT only) or 10 µM ACC, or 0.5 mM linc. Dots represent individual measurements (n = 4), bars are averages. **E**. EIN3 protein accumulation in three-day-old *35S::EIN3-GFP/ein3eil1* seedlings grown in low light on regular growth medium, or medium supplemented with 10 µM ACC or 0.5 mM linc, detected by anti-GFP antibodies. Data is relative to light control and coomassie staining, dots represent biological replicates (n = 3), bars are averages. In A – E different letters mark significant differences, p < 0.05 (A and D: 2-way ANOVA and post-hoc Tukey, B and C: Kruskal-Wallis with post-hoc Dunn, E: 1-way ANOVA with post-hoc Tukey).

Our findings above suggested that the suppression of cotyledon separation in light induced by RS might be a consequence of ET accumulation. To test this end, we first examined the response of the ET-deficient mutant *amiR(acs)* (Tsuchisaka et al., 2009). The conversion of S-adenosylmethionine to ACC is the rate-limiting step during ET production and is catalyzed by ACC-synthase (ACS), which acts as a dimer of various combinations of ACS isoforms (Tsuchisaka et al., 2009). ACC is further oxidized to C_2_H_4_ (ET) by ACC oxidase (ACO). Tsuchisaka *et al*. described that the *amiR(acs)* octuple mutant, deficient in nine ACS isoforms, is unable to form functional dimers and as a consequence produces extremely low levels of ET. We exposed this mutant to lincomycin and found that the synthesis of ET is essential for the lincomycin-induced repression of cotyledon separation in the light (Fig. 2C). Next, we used gas chromatography followed by mass spectrometry (GC-MS) to measure ET emission by seedlings grown on low (1 µM) and higher (10 µM) levels of ACC and compared this to seedlings treated with lincomycin (Fig. 2D). We included *amiR(acs)* and *gun1* mutants as controls for seedlings that lack ACC-synthesis or retrograde signals, respectively. As expected, 10 µM ACC strongly induced ET emission in all three genotypes. Wild-type seedlings treated with 1 µM ACC produced less, but still significant levels of ET compared to control without ACC. Nevertheless, lincomycin did not affect ET emissions in any of the genotypes.

Coherent with the lack of increased ET synthesis, lincomycin did not stabilize EIN3. We used the GFP-tagged, overexpressed EIN3 protein (*p35S::EIN3-GFP / ein3eil1*) to measure protein stability in our experimental set-ups by immuno-blotting. Figure 2E shows that EIN3-GFP is stabilized by 10 µM ACC but not by lincomycin. A smaller protein detected in lincomycin-treated seedlings corresponds to a non-specific product as it was also present in the *ein3eil1* mutant background samples (Fig. S3).

Finally, we tested if the inhibition of cotyledon separation caused by ET is regulated via GUN1-mediated chloroplast signals. We used *gun1* knock-out and *GLK1*-overexpressing seedlings, both with severely reduce sensitivity to lincomycin (Martín et al., 2016). In contrast to lincomycin, ACC could inhibit cotyledon separation in *gun1* and *GLK1-ox*, similar to WT (Fig. S4).

Together, these results indicate that: (1) lincomycin-induced RS and ET inhibit cotyledon separation in light-grown seedlings; (2) chloroplast disruption by lincomycin does not cause changes in ET emissions; (3) GUN1 and GLK1 are not required for the ACC-induced inhibition of cotyledon separation; and (4) ET signaling downstream of EIN3 is required for the lincomycin-mediated inhibition of cotyledon separation in light.

### Lincomycin and ET co-target photomorphogenesis-repressing genes

Because lincomycin and ET regulate similar phenotypes, but RS did not affect ET emissions, we hypothesized that both pathways might co-target a group of photomorphogenesis repressing genes downstream of EIN3. We compared previously published microarray data of low light-grown lincomycin treated seedlings and ACC-treated seedlings and additionally included a dataset of *pif quadruple* (*pifq; pif1pif3pif4pif5*) seedlings grown in darkness (Goda et al., 2008; Leivar et al., 2009; Ruckle et al., 2012). Lincomycin-regulated genes showed strong overlap with either ACC-regulated, or PIF-regulated genes, but only five genes were differentially regulated in all three datasets (Fig. 3A). The violin plot in Fig. 3B represents the fold change of the genes in the three venn-intersections in Fig. 3A. It shows that lincomycin-repressed genes are PIF-repressed in darkness (positive Log_2_(FC) in *pifq*) (317-gene subset), like we previously concluded (Martín et al., 2016). Interestingly, and in contrast to the PIF overlap, it is predominantly the lincomycin-induced gene set that strongly overlaps with ACC-induced genes (95-gene subset). Within this 95-gene subset, 66 genes are co-upregulated by lincomycin and ACC, which is a significant fraction of the complete lincomycin up-regulated gene set. These results are visualized together in the heatmap in Fig. 3C, which presents the same pattern showing that the lincomycin-repressed (negative Log_2_(FC) depicted in yellow)/PIF-repressed (depicted in turquoise) gene set is indeed not regulated by ACC, and that many of the lincomycin-induced genes (positive Log_2_(FC) depicted in turquoise) overlap with the ACC-upregulated set. To explore the role of the EIN3-signalling pathway in the lincomycin-induced RS, we performed additional data comparisons that show no significant enrichment of EIN3 targets among the lincomycin-regulated genes, but does show a significant co-regulation with *ein3eil1* differentially expressed genes (Fig. S5A) (Chang et al., 2013). The lincomycin/ACC co-induced genes do not significantly overlap with PIF-regulated genes (Fig. 3C), as also shown in 3A. Further comparison to light regulated genes (continuous red light versus continuous darkness (Leivar et al., 2009)) shows that, as expected from the lack of PIF-overlap, 94 of the 100 lincomycin/ACC co-regulated genes are not regulated by red light (Fig. S5B, Table S1). We confirmed that the *pifq* mutant has a wild type-like phenotype when grown in lincomycin that can be partly rescued by AgNO3 (Fig. S6), which supports that RS via the ET signaling cascade acts independently of PIF-mediated signaling.

**Figure 3.**
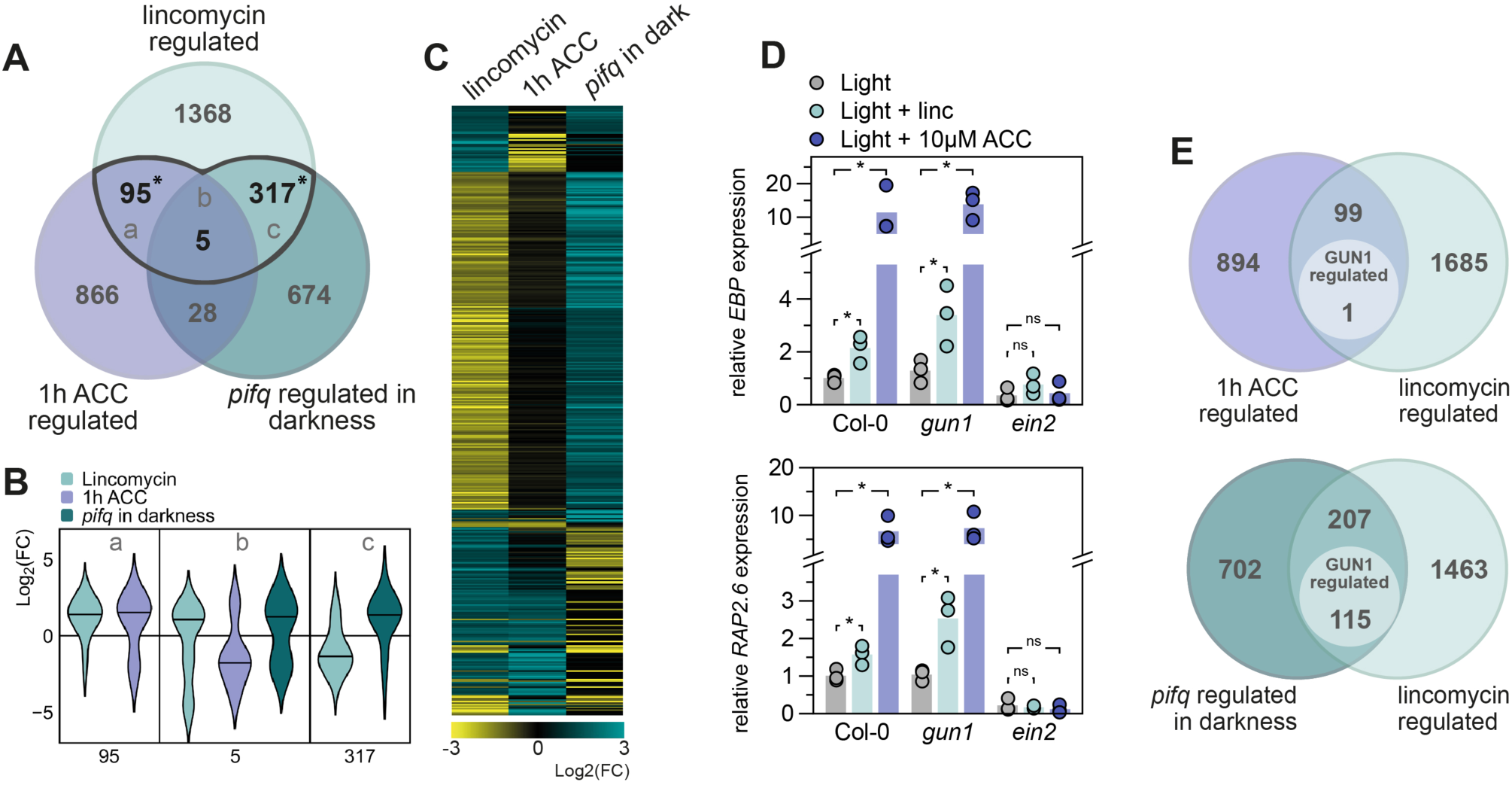
Lincomycin and ACC signaling pathways converge downstream of EIN3. **A**. Overlap of differentially expressed genes (p < 0.05, FC > |1|) in six-day-old low-light grown 0.5 mM linc-treated seedlings, seven-day-old long-day grown 10 µM ACC-treated seedlings, and three-day-old dark-grown *pifq* mutants. Letters (a, b, c) refer to B (Goda et al., 2008; Leivar et al., 2009; Ruckle and Larkin, 2009). Asterisks represent significant overlap (hypergeometric distribution, p < 0.05). **B**. Violin plots representing the Log2(fold change) of the genes differentially expressed by linc and at least one other condition (a, b, c denote the gene sets defined in A.). **C**. Heatmap representing the Log2(fold change) of the genes in B. **D**. Relative expression of *EBP* and *RAP2*.*6* in wild type (WT), *gun1* and *ein2*, grown in low light on regular growth medium, or medium supplemented with 10 µM ACC (left) or 0.5 mM linc (right), measured by RT qPCR. Light control data is similar for left and right graphs. Dots represent biological replicates (n = 3), bars are averages. Significant differences are indicated by asterisks (student T-test, p < 0.05). **E**. Overlap of differentially expressed genes (p < 0.05, FC > |1|). Lincomycin, ACC and *pifq* datasets (as in A), are compared to 5 day-old light-grown and lincomycin-treated (200 ug/mL) *gun1* mutants as compared to wild-type (A and B; GSE5770).

Visual analysis of the lincomycin/ACC up-regulated gene set revealed several transcription factors members of the MYB, bZIP and NAC protein families (Table S1).

### A GUN1-independent retrograde signal targets ethylene signaling components

To investigate whether these lincomycin and ET co-induced genes are possibly regulated by a GUN1-mediated retrograde signal, we analyzed the expression of two genes encoding ethylene response factors: *ETHYLENE-RESPONSIVE ELEMENT BINDING PROTEIN* (*EBP*) and *RELATED TO AP2 6* (*RAP2*.*6*). Both genes were significantly induced by 10 µM ACC and by 0.5 mM lincomycin (Fig. 3D). The induction of *EBP* and *RAP2*.*6* expression by both treatments was lost in the *ein2* mutant, which confirms that the induction of ethylene response factors by lincomycin-induced retrograde signals requires the functional ethylene-response pathway. Interestingly, induced expression by lincomycin was maintained in the *gun1* mutant, which hints towards a GUN1-independent retrograde pathway affecting ET-regulated seedling development.

To further explore this possible GUN1-independence, we extended our analysis and compared the genes co-targeted by lincomycin and ACC (Fig. 3A) to a previously defined list of GUN1-dependent genes. This gene list was derived from light grown, lincomycin-treated seedlings of wild type vs. *gun1* mutants (Koussevitzky et al., 2007). Only 1 out of the 100 lincomycin and ACC co-regulated genes is regulated by GUN1-dependent retrograde signals according to the data of Koussevitzky *et al*. (Fig. 3E). Contrastingly, out of the 322 *pifq* and lincomycin co-targeted genes, 115 (35.7%) are regulated via GUN1 (Fig. 3E). These data strongly suggest that lincomycin induces ethylene-responsive genes independently of GUN1 (Fig. 4). To explore if this GUN1-independent pathway is functionally involved in the closed-cotyledon phenotype, we treated *gun1* mutants with AgNO3 to simultaneously repress the ET signaling pathway. Even though *gun1* seedlings show almost completely separated cotyledons in the presence of lincomycin, the inhibition of ET perception significantly increased the angle between the cotyledons towards control-level (Fig. S6). This supports the presence of a GUN1-independent, ET-dependent, RS cascade which inhibits photomorphogenesis in light-grown seedlings.

**Figure 4.**
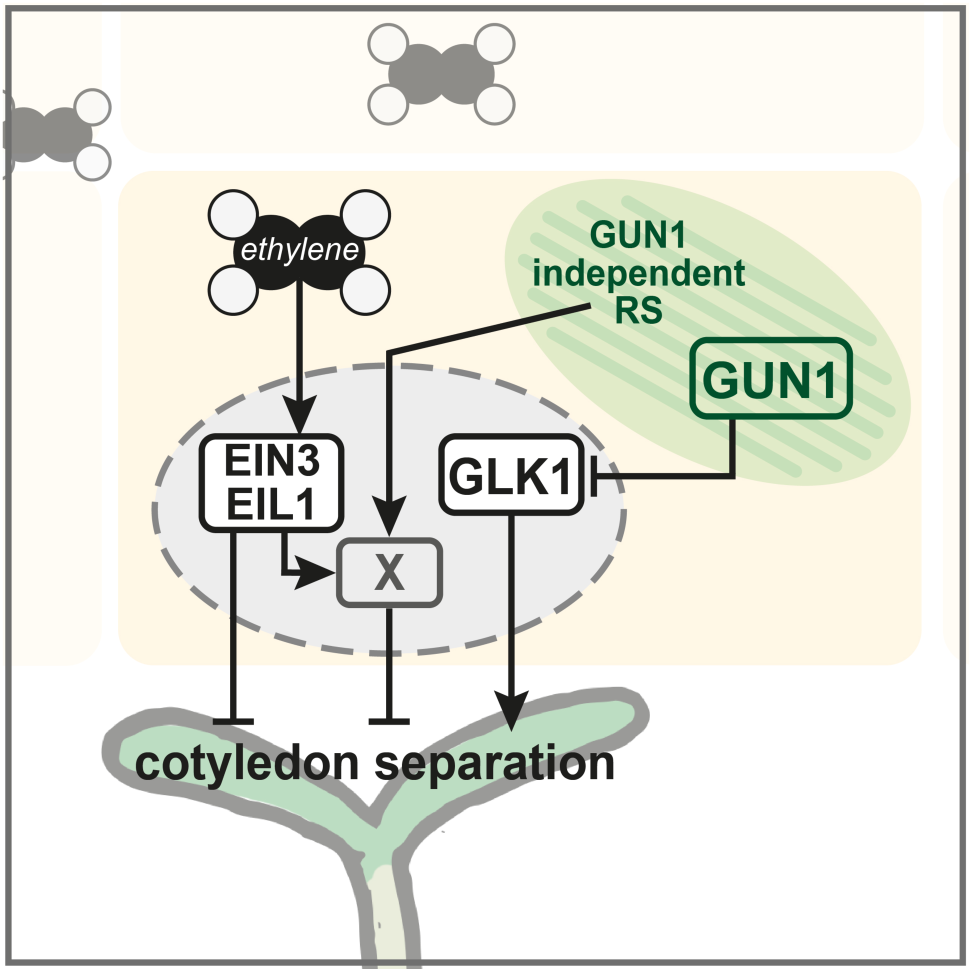
Graphic summary. Ethylene represses cotyledon opening via EIN3 and EIL1. Retrograde signals, induced by lincomycin, repress cotyledon separation via two pathways: a GUN1-dependent repression of photomorphogenesis-inducing factors (e.g. *GLK1*), and a GUN1-independent induction of EIN3 / EIL1 co-targets. “X” represents a repressor of cotyledon separation downstream of EIN3/EIL1 that is induced by ethylene and by chloroplast signals independently of GUN1. X is likely a gene (or set of genes) regulated redundantly by EIN3 and EIL1.

## Discussion

Light strongly drives plant development and promotes photomorphogenesis during seedling establishment. Activated light receptors promote greening and expansion of cotyledons, opening of the apical hook and inhibition of stem growth (Gommers and Monte, 2018). Nevertheless, this process is inhibited when the development of chloroplasts is blocked by drugs such as lincomycin (Ruckle and Larkin, 2009). The retrograde signals released by disrupted chloroplasts target the PIF-repressed phy-induced transcriptome including genes like *GLK1*, to inhibit photomorphogenesis in the light (Martín et al., 2016). Our findings here support a model whereby ethylene (ET) prevents excessive cotyledon separation under continuous light by regulating a set of ET-induced genes downstream of the transcription factor ETHYLENE INSENSITIVE3 (EIN3). In turn, when retrograde signaling (RS) is induced during early photomorphogenesis, inhibition of cotyledon separation by chloroplast signals takes place at least in part by further inducing this same ET signaling pathway. Both signals converge to co-regulate a set of RS/ET-induced genes. The chloroplast signal is yet undefined, but it does not involve GUN1 (Fig 4).

Interestingly, this RS/ET-targeted gene set does not overlap with the RS/PIF-repressed network previously defined in our laboratory (Fig. 3) (Martín et al., 2016). This finding suggests that RS optimizes seedling development in response to chloroplast disruption (caused for example by high light) by targeting at least two distinct gene networks that independently inhibit cotyledon separation: (1) a light-regulated network downstream of the phytochrome/PIF system, and (2) a light-independent ET hormone-regulated pathway downstream of EIN3. Existence of these two separate regulatory pathways is supported by the strong photomorphogenic phenotype of *pifqein3eil1* in darkness compared to *pifq* and *ein3eil1* (Shi et al., 2018). Remarkably, whereas RS antagonizes the light-regulated network by repressing light-induced genes (Martín et al., 2016), our results here suggest that RS further induces an ET-induced network. This dichotomy resulting in the repression or promotion of nuclear gene expression might explain why the chloroplast utilizes two distinct signaling pathways: a GUN1-dependent to repress the light-induced gene network and a GUN1-independent pathway to induce ET-regulated genes. Although retrograde signals other than GUN1 have been described in response to different stresses (Chan et al., 2016), GUN1 is considered a hub factor of RS during chloroplast biogenesis (Mochizuki et al., 1996; León et al., 2013; Hernandez-Verdeja and Strand, 2018; Wu et al., 2018). Our findings here suggest the existence of at least one additional GUN1-independent pathway to optimally adjust seedling development to the light environment.

Cotyledon closing by EIN3 in darkness requires activation of the ethylene signaling pathway by ACC (Shi et al., 2018). We showed genetically and chemically that the inhibition of cotyledon separation by lincomycin also requires functioning ethylene perception and EIN2 and EIN3/EIL1 signaling, but does not enhance ethylene emission. We did not see a stabilization of the EIN3 protein by lincomycin, which is coherent with the lack of induced ET synthesis.

Despite the undetectable stabilization of EIN3-GFP, transcriptional targets of the EIN2 / EIN3 signaling module were up-regulated by lincomycin treatment and ethylene signaling mutants showed reduced lincomycin sensitivity. Different scenarios could explain our results: 1) lincomycin might minimally increase ET synthesis, undetectable in our set-up but sufficient to allow the activation of the signaling cascade downstream of EIN3, 2) lincomycin might stabilize EIN3 transiently during seedling establishment and we might have missed it at the selected time point, or 3) lincomycin might promote an increase in EIN3 activity without affecting EIN3 levels. We consider the first two scenarios most likely, since over-expression of EIN3 could significantly enhance the unseparated cotyledon phenotype compared to the wild type control (Fig. S2A).

The origin of the GUN1-independent retrograde signal that targets the ethylene signaling pathway remains speculative. However, it is interesting to note that the ethylene pathway has been associated with different stress-induced chloroplast signals before. The chloroplast stress-metabolite MEcPP was recently shown to repress ET synthesis and hypocotyl elongation in red light in an auxin-dependent fashion (Jiang et al., 2020). Additionally, reactive oxygen species (ROS) released by mature (green) chloroplasts upon high light stress induce expression of ethylene response factors in the nucleus (Vogel et al., 2014). Lastly, drought and high light stress induce 3’-phosphoadenisone 5’-phosphate (PAP) accumulation in mature chloroplasts that act as RS and inhibit exoribonucleases (XRN’s) such as XRN4, involved in the ethylene signaling pathway as it targets EBF1 and EBF2 mRNA for degradation (Estavillo et al., 2011). Future work will be necessary to address whether these signals are also relevant in young seedlings grown in continuous white light, such as the ones used in our study. We speculate that young seedlings with developing chloroplasts might release biogenic signals via GUN1 as well as ROS and/or PAP to trigger ethylene responses.

To conclude, our results here together with previous work demonstrate that lincomycin represses photomorphogenesis via at least two separate pathways. One requires GUN1-mediated signals and represses PIF-repressed target genes. The other requires GUN1-independent retrograde signals and induces the ethylene response pathway. Future work will explore if this GUN1-independent pathway involves any of the other known retrograde signaling mechanisms, or if it represents a novel and yet undescribed pathway.

## Material and methods

### Plant material and growth conditions

*Arabidopsis thaliana* seeds used here were all described before, including *ein2-5, ein3-1/eil1-1, p35S::EIN2-GFP/ein2-5, p35S::EIN3-GFP/ein3-1/eil1-1, ctr1-1, amiR(acs), gun1-201, p35S::GLK1/glk1glk2* and *pifq*, all in the Columbia-0 wild type background (Kieber et al., 1993; Leivar et al., 2008; Waters et al., 2008; Tsuchisaka et al., 2009; He et al., 2011; Ju et al., 2012; Wen et al., 2012; Xie et al., 2015; Martín et al., 2016). For all experiments, seeds were surface-sterilized and sowed on 0.5MS medium (0.8% plant agar), followed by a four-day stratification treatment before moving to continuous low white light (approx. 1.5 µmol m^-2^ s^-1^, T8 LED tube 4000K, Systion Electronics) at 20°C. Phenotypes, as well as gene, protein and ethylene quantification were analyzed after three days.

### Pharmacological treatments

1-Aminocyclopropane-1-carboxylic acid (ACC; VWR P10007) was dissolved in sterile water (10 mM stock) and added to the growth medium to obtain different final concentrations (0.1, 1, 5, 10, 20, 50 µM. Lincomycin (Sigma-Aldrich L6004) was added directly to the growth medium to obtain 0.5 mM as described before (Martín et al., 2016). Silver nitrate (AgNO_3;_ Sigma-Aldrich 209139) was dissolved in sterile water (1 mM stock) and added to the growth medium in a final concentration of 5 µM. Norflurazon (Sigma-Aldrich 34364) was added directly to the growth medium to obtain a 5 µM solution.

### Phenotype analysis

After three days of growth, seedlings were photographed (Nikon D7000 camera) and pictures were analyzed with Fiji (Schindelin et al., 2012).

### Ethylene measurements

To measure ethylene emissions, seeds were sown in sterilized glass vials (10 ml ‘Headspace Vials’ with screw heads, Restek) which contained 7 ml of growth medium (with or without pharmacological treatment). Each vial contained 120 seeds. Seeds were cold-stratified and germinated as described above. 48 hours after the transfer to light, the air headspace of the vials was flushed with clean air to set a starting point for ethylene accumulation. 24 hours later, measurements were taken. Growth protocol was adapted from (Jeong et al., 2016). 3 ml air was taken from the vial’s headspace and transferred to clean (flushed) Restek vials. Ethylene concentrations were measured by GC-MS with an Agilent 7890A gas chromatograph coupled to a 5975C mass selective detector, as described before (Pereira et al., 2017).

### Transcriptome re-analysis

Published and publicly available micro-array datasets (accession numbers: GSE24517, GSE17159, GSE39384, GSE5770 and GSE21762) were re-analyzed using the GEO2R online tool (NCBI). Differentially expressed genes were selected by 2-fold (Log2 fold > |1|) regulation and adjusted p-value < 0.05. For comparison with ChIP-seq data (Fig. S5A), we have removed the EIN3 target genes that are not represented on the Affimetrix chip which was used for the micro-array experiment.

### Gene expression analysis

To analyze gene expression, 20 seedlings were harvested and pooled per sample, flash-frozen, homogenized and RNA was extracted using the Maxwell total RNA purification kit with DNAse treatment (Promega). cDNA was synthesized using the NZYtech first-strand cDNA synthesis kit with random primers and afterwards treated with RNAse. Quantitative RT-PCR was performed using a Roche Lightcycler 480, SYBR Green mix (Roche) and primers for *EBP* (AT3G16770; 5’-CCCACCAACCAAGTTAACGT-3’ and 5’-GTGGATCTCGAATCTCAGCC-3’), *RAP2*.*6* (AT1G43160; 5’-TGATTACCGGTTCAGCTGTG-3’ and 5’-CTTGTGTGGGTCTCGAATCT-3’). Expression of *PP2A* was analyzed as a reference (AT1G13320; 5’-TATCGGATGACGATTCTTCGT-3’ and 5’-GCTTGGTCGACTATCGGAATG-3’). Relative expression was calculated as 2^(delta-delta CT).

### Protein analysis

Protein extracts were prepared of three-day-old seedlings, pooled per plate for each sample, flash-frozen and homogenized by hand. Nuclear protein was extracted with an extraction buffer as described before (Soy et al., 2014) (buffer: 100 mM MOPS (pH 7.6), 2% SDS, 10% glycerol, 4 mM EDTA, 50 mM Na_2_S_2_O_5_, 2 µgl^-1^ aprotinin, 3 µgl^-1^ leupeptine, 1 µgl^-1^ pepstatin and 2 mM PMSF). Total protein of the samples was quantified using a Protein DC kit (Bio-Rad). β-Mercaptoethanol and loading dye was added to 125 µg of the samples, which were boiled at 95°C for 5 minutes before being loaded on a 7.5% SDS-PAGE gel. Proteins were then transferred to Immobilon-P membrane (Millipore) and EIN3-GFP was detected using an anti-GFP antibody (diluted 1:10.000) (Invitrogen A11122). Anti-rabbit secondary antibody (Sigma NA934) and SuperSignal West Femto chemiluminescence kit (Pierce) were used for protein detection in a Bio-Rad imaging system. The ImageLab program (Bio-rad) was used to quantify band intensity and sizes from blot images, which was compared to a reference from the Coomassie-stained blot.

### Statistical analysis

Multivariate comparisons were done in R and using the online MVApp (Julkowska et al., 2019). First, data was checked for equal variance by Levene’s test and when needed, data was LN-transformed to make variance equal. Multivariate analyses were done by 1 or 2-way ANOVA with a post-hoc Tukey test for pairwise comparisons. For experiments with non-equal variances, the non-parametric Kruskal-Wallis test was applied, with a post-hoc Dunn test or Mann-Whitney for pairwise comparison. Student’s T-test pairwise comparisons were done in Microsoft Excel preceded by an F-test to test for equal variances and LN-transformed when needed. Hypergeometric distribution tests for significant overlap in venn-diagrams were done in R.

## Acknowledgements

We thank Jason Argyris (CRAG, IRTA) for his help to prepare the ethylene measurements, Zeguang Liu, Sjon Hartman and Ronald Pierik (Utrecht University) for sharing seeds of the ethylene signaling mutants, and lab members Arnau Rovira, Nil Veciana, Liu Duan and Ana Couso for their help to harvest materials and support. This work was supported by grants from FEDER / Ministerio de Ciencia, Innovación y Universidades – Agencia Estatal de Investigación (Project References BIO2015-68460-P and PGC2018-099987-B-I00), from the Spanish Ministry of Economy and Competitiveness (FJCI-2016-30876 to C.G.) and from the CERCA Programme / Generalitat de Catalunya (Project Reference 2017SGR-718) to E.M. We acknowledge financial support from the Spanish Ministry of Economy and Competitiveness, through the “Severo Ochoa Programme for Centres of Excellence in R&D” 2016-2019 (SEV-2015-0533)”.

## Supplemental Data

**Figure S1. ACC inhibits photomorphogenesis via EIN2 and EIN3**. Hypocotyl length, cotyledon separation (degrees) and apical hook angle of three-day old wild type Col-0 (WT), *ein2, 35S::EIN2-GFP/ein2* (*EIN2ox*), *ein3eil1* and *35S::EIN3-GFP/ein3eil1* (*EIN3ox*) seedlings grown in continuous low light (approx. 1.5 µmol m^-2^ s^-1^) on normal growth medium, or medium supplemented with different concentrations of ACC. Same data as presented in figure 1 A and B.

**Figure S2. A**. Cotyledon separation (degrees) of three-day-old, light-grown wild type (WT), *ein3-1, eil1-3, ein3eil1* and *35S::EIN3-GFP/ein3eil1* (*EIN3ox*) seedlings on regular growth medium or medium supplemented with 0.5 mM lincomycin (linc). **B**. Cotyledon separation of three-day-old, light-grown wild type seedlings on medium supplemented with 5 µM AgNO_3_, 5 µM Norflurazon (NF), or a combination of both. Different letters indicate significant differences, for A: non-parametric Kruskall wallis, with posthoc Mann-Whitney test; for B: 2-way ANOVA with posthoc Tukey test, p < 0.05.

**Figure S3. Immunoblots for the EIN3-GFP protein**. EIN3 protein accumulation in three-day-old *35S::EIN3-GFP/ein3eil1* seedlings grown in low light on regular growth medium (C), or medium supplemented with 10 µM ACC (A) or 0.5 mM linc (L), detected by anti-GFP antibodies. **A**. First lane are *ein3eil1* seedlings grown with ACC as a negative control. Three biological replicates are represented for all treatments. **B**. Immunoblot of *35S::EIN3-GFP/ein3eil1* and *ein3eil1* seedlings treated with 10 µM ACC (A) or 0.5 mM linc (L) and two blank samples. Protein detected by anti-GFP antibody. n.s. = non-specific band.

**Figure S4. ACC inhibits photomorphogenesis independent of GUN1 and GLK1**. Cotyledon separation (degrees) of three-day-old, light-grown wild type (WT), *gun1* and *35S::GLK1 (GLK1ox)* seedlings on regular growth medium or medium supplemented with 0.5 mM lincomycin (linc) or 10 µM ACC.

**Figure S5. A**. Overlap of differentially expressed genes (p < 0.05, FC > |1|) in six-day-old low-light grown 0.5 mM linc-treated as compared to non-treated wild-type Col-0 seedlings and (left) targets of EIN3 identified by ChIP-seq (Chang et al., 2013) or (right) seven-day-old, long day-grown *ein3eil1* seedlings as compared to wild-type Col-0 (accession no. GSE21762). Asterisk represents significant overlap (hypergeometric distribution, p < 0.05). **B**. Overlap of differentially expressed genes (p < 0.05, FC > |1|) in six-day-old low-light grown 0.5 mM linc-treated seedlings, seven-day-old long-day grown 10 µM ACC-treated seedlings, and three-day-old continuous red light vs. dark-grown seedlings (Goda et al., 2008; Leivar et al., 2009; Ruckle and Larkin, 2009).

**Figure S6. AgNO3 inhibits lincomycin-induced inhibition of cotyledon separation independent of PIFs and GUN1**. Cotyledon separation of three-day-old, light-grown wild type, *pifq* and *gun1* seedlings on medium supplemented with 5 µM AgNO_3_, 0.5 mM linc, or a combination of both. Different letters indicate significant differences, Kruskall wallis with posthoc Mann-Whitney test, p < 0.05.

**Table S1**. List of genes co-regulated by lincomycin and 1h ACC or *pifq* in darkness, which are marked in Fig. 3A and plotted in Fig. 3B, with fold-change and adjusted p-value (Goda et al., 2008; Leivar et al., 2009; Ruckle and Larkin, 2009).

